# Chronic toxicity of three neonicotinoid insecticides and their mixture on two daphniid species: *Daphnia magna* and *Ceriodaphnia dubia*

**DOI:** 10.1101/2021.04.16.440143

**Authors:** Claire Duchet, Chelsea J. Mitchell, Jenifer K. McIntyre, John D. Stark

## Abstract

Neonicotinoid insecticides represent nearly a quarter of the global market and are widely used in agriculture but also for lawn, garden care, and pest control. They are highly water-soluble, persistent in soil, and may enter the aquatic compartment via spray drift, runoff, or leaching, and contribute to downstream aquatic toxicity. Although insects appear to be the most sensitive group to neonicotinoids, other groups, such as crustaceans and birds, may also be affected. Furthermore, most studies focus on single-insecticide exposure and very little is known concerning the impact of neonicotinoid mixtures on aquatic invertebrates. The present study was designed to test potential toxicological effects of an environmentally relevant mixture of imidacloprid, clothianidin, and thiamethoxam on populations of *Ceriodaphnia dubia* and *Daphnia magna* under controlled conditions. Chronic toxicity tests were conducted in the laboratory, and survival and reproduction were measured for both species under exposure to nominal concentrations of imidacloprid (0.256 µg/L), clothianidin (3.11 µg/L), thiamethoxam (1.49 µg/L), and a mixture of the three compounds at the same concentrations of the individual compounds. The neonicotinoids did not affect the survival of *C. dubia* and *D. magna* founders. Reproduction of *C. dubia* was affected only by the mixture. All three individual insecticides as well as the mixture caused a significant reduction in the reproduction of *D. magna*. Our results highlight the complexity of pesticide toxicity and show that traditional toxicological approaches such as acute mortality studies, especially tests with single compounds, can underestimate negative impacts that occur in the environment.

**Highlights:**

- Neonicotinoids are currently the most frequently used insecticides worldwide.
- An environmentally relevant mixture of three neonicotinoids was evaluated on two daphniid species.
- The mixture negatively affected the reproduction of *C. dubia* and *Daphnia magna*.
- Traditional toxicological approaches with single compounds may underestimate the effects occurring in the environment at low concentrations.

## 1. Introduction

Neonicotinoid insecticides represent 25% of the global market, and their use is increasing globally (Borsuah et al. 2020). Clothianidin, imidacloprid, and thiamethoxam are the most commonly used neonicotinoids (Borsuah et al. 2020). They are applied mainly as seed coatings for crops such as corn, soybeans, sunflowers, cotton, and canola (Simon-Delso et al., 2015). Imidacloprid also has a variety of other uses including lawn and garden care, and topical flea medicines (Jeschke et al., 2010).

Neonicotinoid insecticides interfere with the insect nervous system and exhibit very high selectivity for insect nicotinic acetylcholine receptors (*n*AChR), making the neonicotinoids the most important chemical class of insecticides on the global market (Jeschke et al., 2010). The binding to nicotinic acetylcholine receptors is almost irreversible in insects (Jeschke et al., 2010). Therefore, continued exposures of insects to neonicotinoids may lead to a cumulative effect and irreversible blockage of *n*AChRs in their central nervous system (Tennekes and Sanchez-Bayo, 2011), causing paralysis and death. Neonicotinoids have been linked to decreased survival in honeybees (Henry et al., 2012), to reduced colony growth and queen performance in bumble bees (Whitehorn et al., 2012) and sublethal effects on flies with an increase in mating, but a decrease in fecundity (Charpentier et al., 2014).

Although insects appear to be the most sensitive group of animals (Raby et al., 2018a), studies have shown negative effects of neonicotinoids on crustaceans, such as shrimp (Hook et al., 2018), crayfish (Barbee and Stout, 2009), and zooplankton crustaceans such as daphnids and ostracods (Sánchez-Bayo and Goka, 2006). In acute standardized toxicity tests, *Daphnia magna* (Crustacea: Cladocera) appears less sensitive to neonicotinoids among several arthropods tested (LC_50_ (24h) = 7,200 µg/L; Beketov and Liess, 2008). However, the acute LC_50_ (48h) for imidacloprid to *Ceriodaphnia dubia* was reported as 2.1 µg/L, which is lower than usually observed in the literature (Chen et al. 2010).

Neonicotinoids are not intended for use in water bodies, but they enter the aquatic compartment via spray drift, runoff, and leaching (Tišler et al., 2009). In the environment, neonicotinoids are highly soluble in water and persistent in soil, with clothianidin having the longest soil degradation half-life (545 days; Hladik et al., 2014). More and more studies demonstrating the presence of neonicotinoids in water bodies have been recently published. Due to their intensive use in agriculture, mainly as seed coating, neonicotinoids are frequently detected in aquatic ecosystems (Hladik et al,, 2018) at concentrations that may harm aquatic invertebrates. In water samples from wetlands adjacent to agricultural fields in Canada, neonicotinoid concentrations reached a maximum of 3.11 µg/L for clothianidin, 0.256 µg/L for imidacloprid, and 1.49 µg/L for thiamethoxam (Main et al., 2014). These concentrations exceed the ecological thresholds for neonicotinoid concentrations in water to protect aquatic invertebrates, of 0.2 µg/L for short-term (acute) effects and 0.035 µg/L for long-term (chronic) effects (Morrissey et al., 2015). Maximum concentrations of neonicotinoids are often measured in surface runoff samples during storm events following seeding (Chrétien et al., 2017), in water bodies within maize fields during crop season (Schaafsma et al., 2015), or in streams nearby imidacloprid-treated forests after storm events, at concentrations exceeding the USEPA freshwater invertebrate chronic endpoint of 0.01 μg/L (Wiggins et al., 2018). In intensive agricultural regions such as the Midwestern United States, clothianidin, imidacloprid, and thiamethoxam were detected in finished drinking water, which demonstrates their persistence during conventional water treatment (Klarich et al., 2017). Furthermore, neonicotinoids often occur in mixtures at low concentrations in surface waters. In a wide-scale study, Hladik and Kolpin (2016) showed that neonicotinoids were detected in 63% of the streams they monitored across the USA, imidacloprid being the most frequently detected (37%), followed by clothianidin (24%), thiamethoxam (21%), dinotefuran (13%), acetamiprid (3%). They observed that mixtures of multiple neonicotinoid compounds were common; two or more neonicotinoids were detected in 26% of the samples and three or more in 11% of the samples. Unfortunately, few data are available regarding the toxic effects of pesticide mixtures to aquatic organisms, which are more likely to occur in the environment than single pesticide exposures. Moreover, most of the data on neonicotinoid exposure are derived from acute toxicity tests. Although acute lethal toxicity of neonicotinoids is a concern, sublethal endpoints under chronic exposure are crucial for assessing the survival of invertebrate populations exposed to neonicotinoids in environmentally realistic conditions (Cavallaro et al., 2017). Therefore, in the study presented here, we evaluated the toxicity (lethal and sublethal) of imidacloprid, clothianidin, and thiamethoxam, separately and as a mixture at environmentally relevant concentrations, to two cladoceran species, *Daphnia magna* and *Ceriodaphnia dubia*.

## 2. Material and Methods

### 2.1. Insecticides

We evaluated commercial formulations of each insecticide rather than technical products because these are the products applied for pest control. Imidacloprid was applied as the flowable formulation Admire®Pro Systemic Protectant (42.8% active ingredient (AI); CAS # 138261-41-3, Bayer CropScience, Leverkusen, Germany). Clothianidin was applied as Belay® Insecticide (23% AI, CAS # 210880-92-5, Valent U.S.A. Corporation, Walnut Creek, CA, USA). Thiamethoxam was applied as Platinum® 75SG (75% AI, CAS # 153719-23-4, Syngenta Crop Protection, Greensboro, NC). Nominal insecticide concentrations were prepared from serial dilutions in deionized water of freshly prepared stock solutions (10.24 µg/L AI for imidacloprid; 124.4 µg/L AI for clothianidin; 59.6 µg/L AI for thiamethoxam). Stocks were subsequently diluted to achieve the environmentally relevant target test concentrations of 0.256 µg/L AI imidacloprid, 3.11 µg/L AI clothianidin and 1.49 µg/L AI thiamethoxam, and a mixture of the three compounds at the same nominal concentrations. Those concentrations were chosen according to environmental concentrations measured in water bodies adjacent to agricultural field (Main et al., 2014). The concentrated stock solutions of the insecticides prepared at the beginning of the experiment were left aging during the experiment, to mimic chemical degradation. The stock solutions were stored under the same environmental conditions as the test organisms (same temperature and light regimen).

### 2.2. Toxicity tests

Assays were performed using the 4th brood offspring of *C. dubia* and *D. magna* cultured at the Washington State University Puyallup Research and Extension Center (Puyallup, WA, USA) according to a method modified from a USEPA protocol (EPA, 2002). Each species was reared in 1 L glass aquaria filled with culture water (Millipore Milli-Q® filtered water to which was added nutrients according to EPA, 2002) and maintained at 24 ± 1°C in a light:dark regimen of 16:8. Daphniids were fed three times a week with a suspension of 1:1 mixture of yeast-cereal leaves-trout chow (YCT) and the algal species *Raphidocelis subcapitata* (previously *Pseudokirchneriella subcapitata*; Aquatic Research Organisms, Hampton, NH) while renewing the medium water.

Tests were conducted in 50 ml glass beakers containing 20 ml or 40 ml of exposure medium for *C. dubia* and *D. magna*, respectively; 0.5 mL of insecticide stock solution for *C. dubia* or 1 ml for *D. magna*, was added to reconstituted/medium water (see EPA, 2002) containing 0.1 mL of YCT: algae suspension. Single-pulse exposures were performed on neonates (<24 h old) of *C. dubia* or *D. magna* (1 individual per beaker, 10 replicates per treatment, respectively, see EPA procedure for static-renewal tests). The duration of the test was 8 days for *C. dubia* and 21 days for *D. magna* to allow time for at least 3 broods to be produced under the conditions of our study. Dead individuals and newborns were counted and removed when present, every day for *C. dubia* and every two days for *D. magna*, to measure survival and reproduction. Then, surviving founders were transferred to newly made water medium, fed with YCT: algae mixture and exposed to the ageing concentrated stock solutions of the insecticides.

### 2.3. Chemical analyses

Samples from the stock solution of each treatment (125 mL) were collected in polyethylene bottles (Nurnberg Scientific, Tualatin, OR, USA) after each water change during the test, and were stored at −20°C until analysis. Samples were analyzed using liquid chromatography – tandem mass spectrometry (LC-MS/MS), to confirm the nominal concentrations at the beginning of the experiments and to follow the degradation of the insecticides over time (Supplementary Material S1 and T1).

### 2.4. Statistical analysis

Survival of founders (females used to start populations) were analyzed with a Fisher’s exact test to compare the survival between each treatment and the control, as recommended by the USEPA protocol (EPA, 2002). The reproduction of the founders (number of offspring per females) was compared among the treatments and the control with a one-way ANOVA (analysis of variance). Data were square root-transformed to satisfy parametric assumptions prior to analysis. When significant, ANOVA was followed by a Dunnett’s post hoc test (α = 0.05). All tests were performed using RStudio for Windows Version 1.1.453 (RStudio Team 2015). Significance was accepted at α = 0.05 for all tests.

## 3. Results

### 3.1. Chemical analysis

Measured concentrations and percentage differences between measured and nominal concentrations of each pesticide in the stock solutions for each treatment are summarized Table 1. The % difference between nominal and measured concentrations at day 0 was used to correct the nominal concentrations the daphniids were exposed to during the experiment.

**Table 1.** Measured insecticide concentrations (µg/L) in the stock solutions during the experiment and comparison (% difference) with the nominal concentration at the beginning of the experiment (day 0). IMI: imidacloprid; CLO: clothianidin; THM: thiamethoxam; only: single compound solution; mixture: mixture solution.

| Compound | Nominal concentration ($\mu\text{g/L}$) | Measured concentrations ($\mu\text{g/L}$) | | | | % difference | *n* |
| --- | --- | --- | --- | --- | --- | --- | --- |
|  |  | Day 0 | Day 2 | Day 8 | Day 21 |  |  |
| IMI - only | 10.24 | 3.6 | 3.8 | 2.7 | 0.91 | 65 | 1 |
| CLO - only | 124.40 | 100 | 94 | 76 | 37 | 20 | 1 |
| THM - only | 59.60 | 57 | 55 | 50 | 41 | 4 | 1 |
| IMI - mixture | 10.24 | 4 | 3,7 | 2,6 | 1 | 61 | 1 |
| CLO - mixture | 124.40 | 100 | 92 | 74 | 42 | 20 | 1 |
| THM - mixture | 59.60 | 60 | 56 | 47 | 32 | -1 | 1 |

### 3.2. Toxicity testing: Effect on survival and reproduction

Control survival was 90% at the end of both bioassays for both *C. dubia* and *D. magna*, and mean control reproduction was 17 ± 3 for *C. dubia*, and 71 ± 6 for *D. magna*. No ephippia (resting eggs) were seen in any replicate. Therefore, both bioassays passed the test validity criteria which are control survival ≥ 80% and reproduction of > 15 neonates per *C. dubia* female and > 60 neonates per *D. magna* female (EPA 2002).

The survival of the *C. dubia* and *D. magna* founders exposed to individual the compounds or the mixture was not statistically different from control (Fisher’s exact test, *p* > 0.05; Table 2).

**Table 2.**
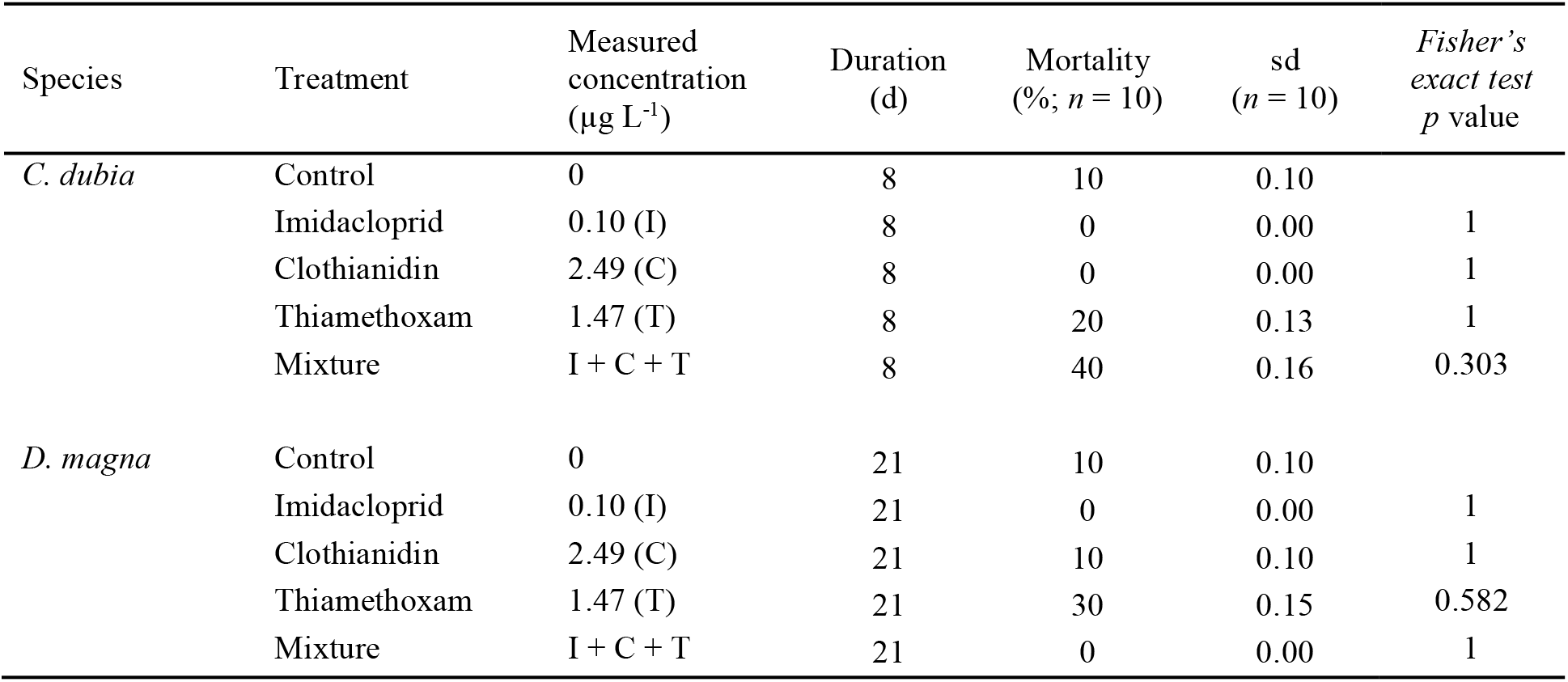
Survival of *Ceriodaphnia dubia* and *Daphnia magna* (n=10 each) exposed to each of the three neonicotinoids and the mixture.

| Species | Treatment | Measured concentration ($\mu\text{g L}^{-1}$) | Duration (d) | Mortality (%; $n = 10$) | sd ($n = 10$) | Fisher's exact test $p$ value |
| --- | --- | --- | --- | --- | --- | --- |
| *C. dubia* | Control | 0 | 8 | 10 | 0.10 |  |
|  | Imidacloprid | 0.10 (I) | 8 | 0 | 0.00 | 1 |
|  | Clothianidin | 2.49 (C) | 8 | 0 | 0.00 | 1 |
|  | Thiamethoxam | 1.47 (T) | 8 | 20 | 0.13 | 1 |
|  | Mixture | I + C + T | 8 | 40 | 0.16 | 0.303 |
| *D. magna* | Control | 0 | 21 | 10 | 0.10 |  |
|  | Imidacloprid | 0.10 (I) | 21 | 0 | 0.00 | 1 |
|  | Clothianidin | 2.49 (C) | 21 | 10 | 0.10 | 1 |
|  | Thiamethoxam | 1.47 (T) | 21 | 30 | 0.15 | 0.582 |
|  | Mixture | I + C + T | 21 | 0 | 0.00 | 1 |

Only the mixture of these insecticides had a negative effect on reproduction of *C. dubia*, (Dunnett’s post hoc test following ANOVA, *p* = 0.019, Fig. 1), whereas all three single neonicotinoid compounds and their mixture negatively affected the reproduction of *D. magna* (Dunnett’s post hoc test following ANOVA, *p* < 0.05, Fig. 2).

**Fig.1.**
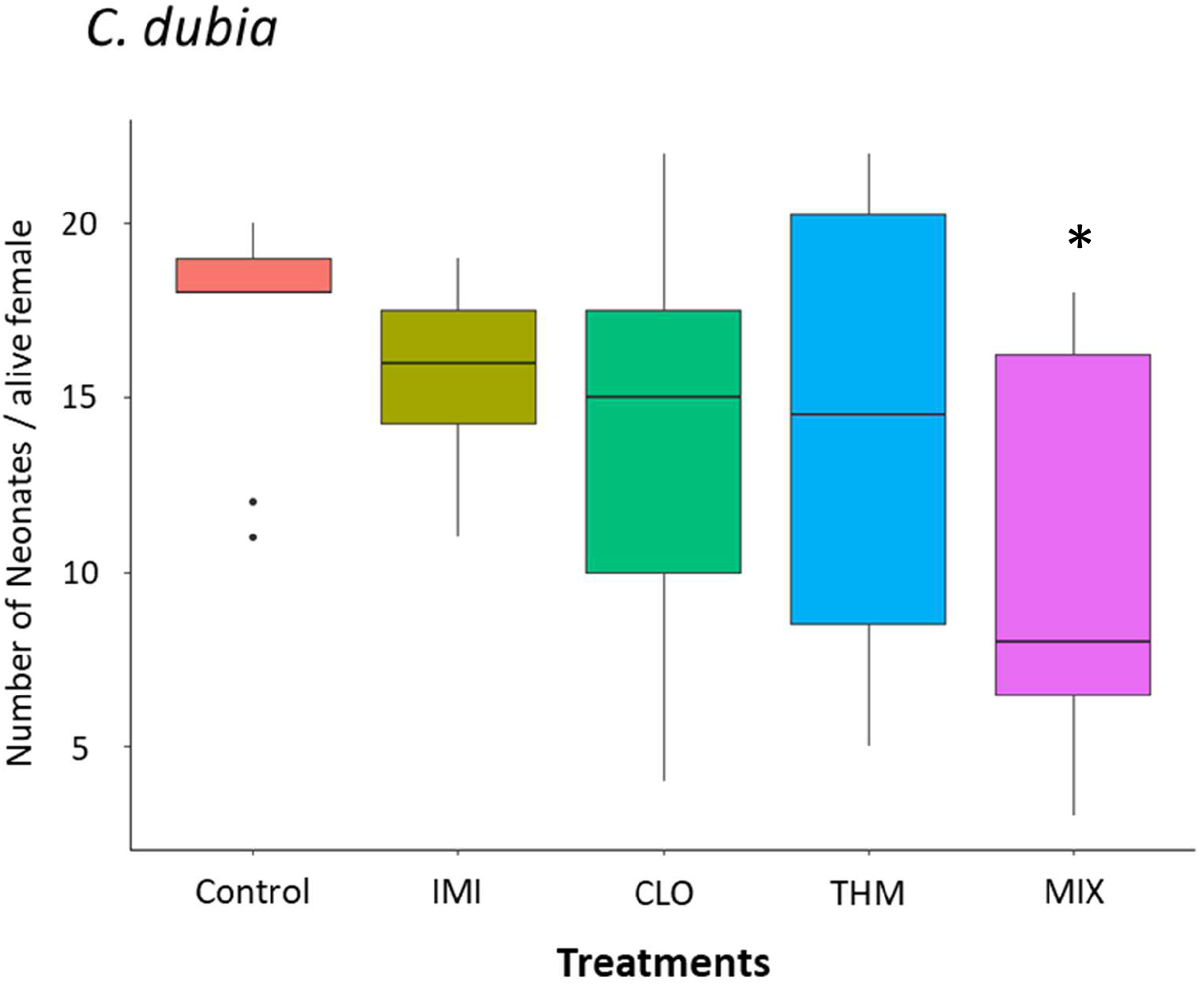
Effects of neonicotinoids on *C. dubia* reproduction (number of neonates per female produced during the test, mean ± SE; *n* = 10 for each treatment). Boxplots represent the median (center line), 1^st^ and 3^rd^ quartiles (hinges), largest value no greater than 1.5 times the interquartile range (IQR) from the hinge (upper whisker), and the smallest value no less than 1.5 times the IQR from the hinge (lower whisker). Dots represent outliers that fall beyond 1.5 times the IQRA. Control: control; IMI: imidacloprid; CLO: clothianidin; THM: thiamethoxam; MIX: mixture of the 3 insecticides. *, **: Significant differences between the treatment and control (Dunnett’s test following ANOVA; *: p < 0.05; **: p < 0.01).

**Fig.2.**
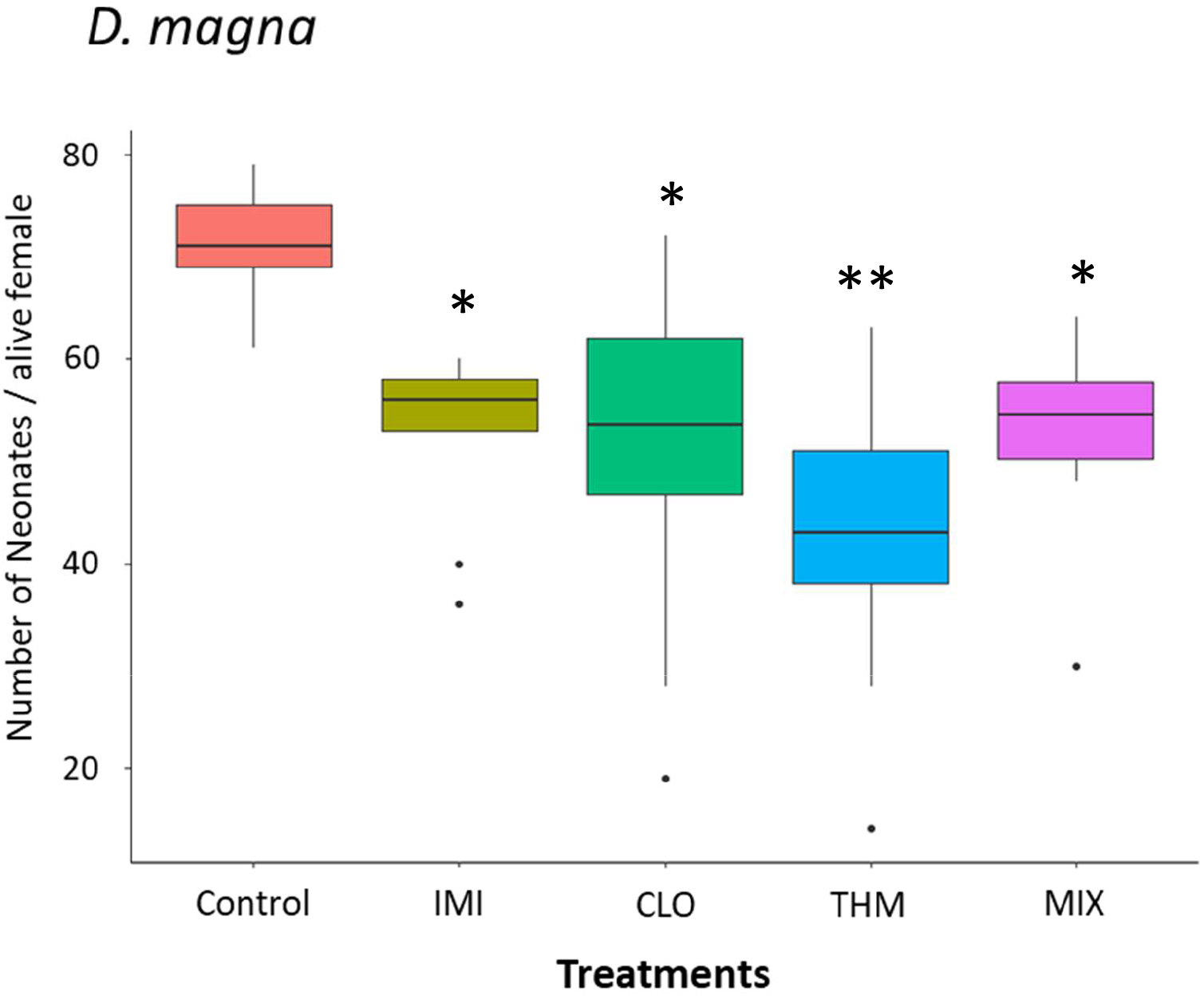
Effects of neonicotinoids on *D. magna* reproduction (number of neonates per female produced during the test, mean ± SE; *n* = 10 for each treatment). Boxplots represent the median (center line), 1^st^ and 3^rd^ quartiles (hinges), largest value no greater than 1.5 times the interquartile range (IQR) from the hinge (upper whisker), and the smallest value no less than 1.5 times the IQR from the hinge (lower whisker). Dots represent outliers that fall beyond 1.5 times the IQRA. Control: control; IMI: imidacloprid; CLO: clothianidin; THM: thiamethoxam; MIX: mixture of the 3 insecticides. *, **: Significant differences between the treatment and control (Dunnett’s test following ANOVA; *: p < 0.05; **: p < 0.01).

## 4. Discussion

Results of this study showed that determining the potential impact of environmentally realistic concentrations of neonicotinoid insecticides on daphniid populations is not straight-forward. Exposure of *C. dubia* and *D. magna* to three individual neonicotinoids and to the mixture did not cause significantly different mortality in founders compared to the controls across the test period. However, the survival of *C. dubia* exposed to the mixture and the survival of *D. magna* exposed to thiamethoxam were lower than control. The mortality, which appeared after 5 days for *C. dubia* exposed to the mixture and after 12 days for *D. magna* exposed to thiamethoxam, may be ecologically significant despite the fact that no significant differences were detected with the Fisher’s exact test. Therefore, the chronic toxicity test, as presented by the EPA, is not appropriate to evaluate delayed effects on the survival due to a low number of founders used in the experiment (10 individual per treatment) and due to the lack of power of the statistic test of the Fisher’s exact test. This needs further consideration to draw a clear conclusion, and delayed mortality induced by the thiamethoxam and the mixture on the daphniids, should be explored further. Indeed, delayed lethal and sublethal effects of neonicotinoids on *D*. magna, have been observed 4 to 12 days following exposure, by Beketov and Liess (2008). By examining longer exposures with a higher number of replicates and founders, it may be possible to determine that neonicotinoids at low environmental concentrations, could have a negative effect on survival of both daphniid species evaluated in this study. This underlines a time-dependent effect, observed whenever a toxicant permanently binds to a receptor in the body (Tennekes and Sanchez-Bayo, 2011).

Exposure to each neonicotinoid separately reduced offspring production in *D. magna*. These results tell a cautionary tale about estimating pesticide effects using only one endpoint of effect (mortality) and only one species. Thiamethoxam had a stronger negative effect on the reproduction of *D. magna* than imidacloprid and clothianidin alone or the mixture. This result is not consistent with the published literature as imidacloprid and clothianidin are often shown to be more toxic than thiamethoxam on freshwater invertebrate, such as *C. dubia* (Raby et al. 2018a), *Hyalella azteca* (Bartlett et al. 2019) or *Chironomus dilutus* (Cavallaro et al. 2017), and this may need to be explored further. Unlike *D. magna*, exposure to individual neonicotinoid insecticides did not significantly affect reproduction of *C. dubia*. However, the reproduction of *C. dubia* founders exposed to the neonicotinoids was on average lower than the controls and the high variability between the individuals may have reduced the power analysis. In a study comparing the chronic sensitivity of *D. magna* and *C. dubia* to imidacloprid, Raby et al. (2018b) showed that *C. dubia* was more sensitive to the neonicotinoid than *D. magna*. In the same study, imidacloprid induced the highest toxicity to *C. dubia*, while thiamethoxam did not induced any negative effects on the survival nor on the reproduction. In our study, thiamethoxam induced a lower survival of *D. magna* when applied as a single compound. However, all three single compounds of neonicotinoids reduced the reproduction of *D. magna* and, and when applied as a mixture, they affected the reproduction of both species, at low concentrations.

Cladocerans are by far the most tolerant taxa among arthropods to neonicotinoid insecticides, with 2 to 3 orders of magnitude lower sensitivity than any other taxa among the arthropods (Morrissey et al., 2015; Raby et al., 2018a). Indeed, neonicotinoids are more toxic to aquatic insects, particularly to the larval stage of ephemeropterans, trichopterans and dipterans, which have been shown to exhibit short-term lethal effects at concentrations below 1 μg/L (Morrissey et al., 2015; Maloney et al., 2018b). Conflicting results between our study and previous studies might be explained by the type of products used in the assays. Pesticide formulations applied for plant protection contain various additives, besides the active ingredient. Those additives have long been considered as inert/inactive ingredients and only require simplified risk assessment compared to the active ingredients (European Council, 2004). However, studies have increasingly found that these additives may play a role in modifying toxicity. Takács et al. (2017) estimated that a formulated product of clothianidin was 46 times more toxic to *D. magna* than clothianidin alone. Jemec et al. (2007) found that the formulated product Confidor SL200 (*a.i*. imidacloprid) was four times more toxic to *D. magna* than the active ingredient alone, explaining the difference of toxicity as a result of synergism between solvents and imidacloprid. Similar differential toxicity of formulated products compared to their active ingredients have been demonstrated for other formulated pesticide products such as glyphosate (Mesnage et al., 2013; Székács et al,, 2014). Results of these studies indicate that appropriate environmental risk assessment of formulations used in agriculture should include the toxicological evaluation of surfactants and other additives.

Most studies on neonicotinoid toxicity assessed acute toxicity to one pesticide at a time (Whitfield-Aslund et al., 2017) and studies assessing the sublethal effects of a neonicotinoid mixture on aquatic invertebrates, and more specifically on reproduction, are scarce. Imidacloprid has been by far the most studied neonicotinoid (66% of 214 toxicity tests reviewed by Morrissey et al., 2015). Where mixture toxicity studies exist, they usually focus on gaining a preliminary understanding of toxicity under acute, high concentration exposure scenarios (Loureiro et al., 2010; Pavlaki et al., 2011). This specific scenario applies when aquatic invertebrates are exposed to pulses of high pesticide concentrations associated with spray and rain events (Cedergreen and Rasmussen, 2017), but can be unrealistic, since in the environment aquatic invertebrates are mainly exposed to chronic exposure of low pesticides concentrations. Despite a measured concentration of imidacloprid noticeably lower than the nominal concentration, it is consistent with concentrations measured in the environment. Indeed, in their study, Main et al. (2014) measured maximum concentrations of imidacloprid of 0.097 µg/L (± 0.138 µg/L) in average, in Prairie wetlands nearby agriculture fields. Recent studies have focused on the effects of neonicotinoid mixture on aquatic invertebrates (Loureiro et al., 2010; Pavlaki et al., 2011; Maloney et al., 2018a) and have underscored the complexity of extrapolated lab-derived mixture models to predict the effects on field populations (Maloney et al., 2018a). We observed a greater negative effect of the neonicotinoid mixture on *Ceriodaphnia dubia* than after exposure to each compound alone. Maloney et al.(2018a) reported that binary and ternary mixtures of imidacloprid, clothianidin, and thiamethoxam had greater than additive effects on the freshwater midge *Chironomus dilutus*. However, our study was not designed to assess additive and synergetic effects of neonicotinoid mixtures on daphnids and this should be addressed in further experiments.

Chronic exposures can sometimes result in much higher mortality levels in populations than predicted by acute, short-term exposures, due to delayed lethal effects and because sublethal effects may affect several population traits, most notably by decreasing fecundity (Forbes and Calow, 1999; Stark and Banks, 2003). Our study showed that mixtures of neonicotinoids affect the reproduction of *C. dubia* and the reproduction of *D. magna*, which can have dramatic effects on the population. Reduced population growth is particularly critical to daphniids, which typically require a high rate of population growth to persist through periods of high mortality from predators (Dodson and Hanazato, 1995). This may lead to significant perturbations at the ecosystem level, due to their key position in aquatic food webs, as they control water quality through selective consumption of algae (Luecke et al., 1992) while providing a major dietary component for several fish species (Post et al., 1992).

## 5. Conclusion

Our results showed a negative effect of a mixture of neonicotinoid insecticides on *C. dubia* and *D. magna* reproduction. This highlights the complexity of evaluating pesticide toxicity and shows that traditional toxicological approaches such as acute mortality studies, especially tests with single compounds, may underestimate impacts that occur in the environment at low pesticide concentrations.

## 6. Author Contribution

CD proposed and designed the study. CD performed the lab work, data gathering, data analysis with the assistance of CJM and carried out the manuscript preparation. All authors contributed critically to the drafts and gave their final approval to the manuscript submitted for publication.

