## Supplementary material S1 for "Chronic toxicity of three neonicotinoid insecticides and their mixture on two daphniid species: *Daphnia magna* and *Ceriodaphnia dubia*"

---

**S.1.** Chemical analyses

Samples collected for the analysis of the neonicotinoid concentrations were stored 3 months at -20ºC before analysis by the Pacific Agricultural Laboratory (Sherwood, OR, USA).

All water samples were analyzed with liquid chromatography/tandem quadrupole mass spectroscopy using a direct injection technique. Samples were prepared for analysis by adding 0.5mL of sample to 0.5mL methanol in an autosampler vial. Additional dilutions in methanol were performed as necessary to remain within the calibration range of the instrument.

An eight-point calibration curve for each target compounds was created ranging from 0.1 ng/mL to 20 ng/mL.

Method detection limits were 0.020 µg L-1 for imidacloprid and thiamethoxam, and 2 µg L-1 for clothianidin.

**Table S1.** LC-MS/MS Mass Transitions

| Compound | Precursor Ion | Product Ion |
|---|---|---|
| Clothianidin | 250.0 | 131.9 |
| Clothianidin | 250.0 | 169.0 |
| Clothianidin | 250.0 | 112.9 |
| Imidacloprid | 256.0 | 208.9 |
| Imidacloprid | 256.0 | 175.0 |
| Thiamethoxam | 292.0 | 211.1 |
| Thiamethoxam | 292.0 | 181.1 |
| Thiamethoxam | 292.0 | 132.0 |
